# Protected areas show substantial and increasing risk of wildfire globally

**DOI:** 10.1101/2025.09.01.673466

**Authors:** Víctor Resco de Dios, Àngel Cunill Camprubí, Ahimsa Campos-Arceiz, Hamish Clarke, Yingpeng He, Obey Zveushe, Rut Domènech, Han Ying, Yinan Yao

## Abstract

Protected area coverage is set to expand in response to climate change and the biodiversity crisis, but we lack assessments of wildfire incidence in protected areas. Here we quantify bigeographical variation in global patterns of burned area in protected areas. During the twenty-first century, wildfires have burned 2 billion hectares of protected areas – an area the size of Russia and India combined – and, while protected areas only cover 19.2% of semi-natural ecosystems, they concentrate 28.5% of the area burned annually. Wildfire in protected areas increased significantly between 2001-2024 (+0.46% yr^-1^), even after taking into account increases in protected area (+0.27% yr^-1^), pointing to a disproportional impact of fire on protected areas under increasingly severe fire weather. This pattern showed marked variation across biomes, with the largest disproportionate increases occurring in fire-prone biomes (e.g. Mediterranean and dry tropical forests, tropical grasslands and xeric shrublands). There were important exceptions to this general trend, and protected area fire was lower than expected in biomes where fire activity is naturally limited by moisture (e.g. tropical rainforests or montane grasslands). Wildfires are important for the health of many ecosystems, and such values of burned area will not always mean a negative outcome. Amidst concerted efforts to expand protected area coverage such as the Global Biodiversity Framework, our results highlight the need for new management strategies that address the globally increasing impacts of burned area across protected areas under unabated climate change.

## Introduction

Anthropogenic activities are catalysing a global polycrisis where multiple stressors interact synergistically and aggravate environmental impacts (Homer-Dixon et al., 2021). Climate change, for instance, is exacerbating the biodiversity crisis by degrading habitats and driving large-scale species adaptation, migration, or—in the worst case scenario—extinction (IPBES, 2019). A less frequently considered feedback between biodiversity and climate change lies in the surge in wildfire activity experienced in recent years, where the frequency and severity of extreme fires has increased (despite declines in overall area burned globally) (Fernández-García and Alonso-González, 2023). Protected areas (PAs) are considered a cornerstone for biodiversity conservation (Langhammer et al., 2024) and ensuring effective nature protection under increasing wildfire incidence is a key challenge for the 21st century.

There is substantial uncertainty as to whether protected areas show a proportionally higher or lower fire activity, relative to unprotected areas (Doherty et al., 2024). This is because assessments of burned area within PAs have so far been performed locally or regionally, and the results vary widely across regions (Nelson and Chomitz, 2011; Rodrigues et al., 2023). We still lack a robust global quantification of how fire impacts PAs as, under the Kunming-Montreal Global Biodiversity Framework (GBF), as PA coverage should increase up to 30% of the land by 2030. Quantifying whether protected area fire is changing will be crucial to the successful implementation of the GBF and other conservation efforts.

Here we quantify, for the first time, variation in the biogeographic patterns of burned area within PAs across biomes, and how they have changed between 2001 and 2024. Also, given that PAs’ coverage has increased steadily during the 21^st^ century, we addressed the critical question of whether the incidence of protected area fire has been increasing proportionally, at the same pace as the expansion of PAs, or whether the increase in PAs has been accompanied by disproportionate increases (overproportion) or decreases in burned area within PAs. The goal of this short note was not to address whether the establishment of forest reserves has led to changes in the fire regime, but to quantify fire incidence in protected areas globally.

## Materials and Methods

### Global Maps of Protected Areas

To quantify the incidence of wildfire inside PAs, we focused our analysis on natural and semi-natural “burnable” vegetation (forests, shrublands, grasslands, and savannas) and we overlaid annual burned area from the Moderate Resolution Imaging Spectroradiometer (MODIS) Burned Area product (MCD64A1) collection 6.1, along with maps of PAs coverage collated from the World Database of Protected Areas (UNEP-WCMC and IUCN, 2023) and other sources, and the definition of biomes from Dinerstein et al. (Dinerstein et al., 2017).

We used the WDPA database (January 2025 version) (UNEP-WCMC and IUCN, 2023) as the main data source on global protected areas. We calculated PA coverage following the guidelines provided by the UN Environment Programme World Conservation Monitoring Centre (UNEP-WCMC). We only considered terrestrial PAs, and the terrestrial part of marine protected areas. Following current practice (Jones et al., 2018; Maxwell et al., 2020; UNEP-WCMC and IUCN, 2023), PAs designated as “Proposed” and “Not reported” were excluded as their implementation is not yet finalized. UNESCO Man and Biosphere (MAB) Reserves were similarly excluded following (UNEP-WCMC and IUCN, 2023) because MAB Core areas are already accounted in WDPA.

When PA establishment dates were missing, we followed previously published approaches (Butchart et al., 2015; Jones et al., 2018; Maxwell et al., 2020) by randomly selecting a year (with replacement) from all PAs within the same country with a known establishment date. For countries with fewer than five protected areas with known establishment date, we randomly selected a year from all terrestrial protected areas with a known date. The random assignment was repeated 1,000 times to identify the median year of establishment, which was then assigned to each protected area with unknown establishment dates.

Some countries apply data restrictions to the WDPA and data could not be accessed to for China, India, Turkey, Eritrea, Western Sahara and the regions of Jammu and Kashmir and Azad Kashmir. Regarding China, we used additional data from the China Nature Reserve Specimen Resource Sharing Platform (http://www.papc.cn/html/folder/946895-1.htm, June 2024). The total coverage was of 1,721,008 km^2^ (∼18% of total land area), which coincides with PA coverage according to other studies (Li and Pimm, 2020). Data on protected areas for India was obtained from the Wildlife Institute of India (https://wii.gov.in, June 2024), and it was adapted to the WDPA structure. We searched the spatial information of Indian PAs with Open Street Map using an OverPass query (https://overpass-turbo.eu, June 2024). We obtained a total of 691 PAs, covering an area of 149,443 km^2^, also in line with other publications.

### Burned Area and Land Cover maps

Regarding spatial burned area estimates, we used the MODIS Burned Area product (MCD64A1) Collection 6.1 in geographic coordinates format. MODIS BA provides global information on burned area in a 500 m grid and monthly time steps (Giglio et al., 2021). We created monthly binary layers by setting a value of 1 to burned pixels, and aggregating them at yearly steps. Aggregated rasters were then multiplied for a cell-size raster layer calculated in spherical coordinates. This corrected for pixels burning more than once in a year.

We used MODIS land cover (LC) type product (MCD12Q1) collection 6.1 (Friedl and Sulla-Menashe, n.d.) to filter burned area and protected areas to specific LC types, following the IGBP scheme. More specifically, we only studied fire in semi-natural vegetation which includes forests (evergreen needleleaf forests, evergreen broadleaf forests, deciduous needleleaf forests and deciduous broadleaf forests), shrublands (open and closed), savannas (woody savannas and savannas) and grasslands. This mask was necessary to avoid artefacts related to a higher presence of non-flammable crops outside protected areas, which could artificially increase the impact of fires within protected areas. This mask was applied considering the conditions of the year before a given fire. That is, the analysis of a fire occurring in 2020 would be based on the value of the mask released at the end of 2019.

We used MODIS LC layers as a template grid to rasterize vector data, as in previous publications (Dobrowski et al., 2021). Country and continent boundary layers were converted to binary rasters at MODIS resolution (∼ 500m) after applying the rule of the 50% cell cover. PA polygons were similarly dissolved by year. Each pixel was then associated with a specific continent/region and characterized as either protected or unprotected following previous publications (Dobrowski et al., 2021), and the LC mask was then applied. MODIS resolution was then aggregated at a 0.25º × 0.25º grid for spatial analyses at global scale.

### Statistical analyses

Our primary response variables were the cover of protected areas across biomes (in absolute terms and also in percent), and protected area fire (the percent of burned area within each protected area biome, %PAF). The absolute cover of PAs was estimated by adding the area of all PAs within a biome, after applying the land use mask. The percent cover of PAs (%PA) within a biome was calculated from the ratio between the absolute cover of PAs and the total area (protected and unprotected) of that biome (after applying the land mask):

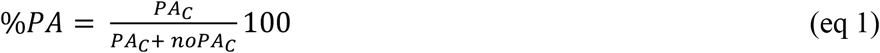

where PA_C_ and noPA_C_ represent the cover of protected and unprotected areas, respectively. %PAF was estimated from the fraction of the total burned area that occurred within PAs (after applying the land cover mask):

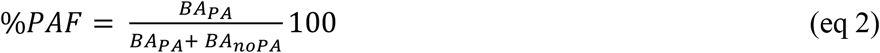

where BA_PA_ and BA_noPA_ indicate the sum of burned area across protected and unprotected areas, respectively.

Temporal trends across of the response variables during study period were estimated using the Mann-Kendall (MK) test and the Sen’s slope, which are common techniques in time series analyses(Bousfield et al., 2023). The former is a non-parametric test, based on ranks, that is used to detect the presence of a monotonic tendency in a time series (Kendall, 1955; Mann, 1945). The latter quantifies the overall slope based on the median of all slopes calculated between each pair of points in the time series.

In order to quantify whether PAs were burning out of proportion, we compared %PA with %PAF. If protected areas burn preferentially, then we should observe that %PAF is significantly larger than %PA. For example, if protected areas occupy 10% of the forest area within a biome (%PA = 10%), but 50% of the burned area occurs in protected areas (%PAF = 50%), then fires would be disproportionately affecting protected areas (%PAF>%PA). If there is no effect of protection on burned area, we expected that %PA would not be significantly different from %PAF (%PAF = %PA). Finally, if protected areas are less affected by fire than the rest of the landscape, then protected area fire should be smaller than the percent of land occupied by protected areas (%PAF < %PA). We assessed for statistical differences between annual values of %PAF and %PA using the MK test and Sen’s slope with D (the difference between slopes). The first test informs on whether both slopes are significantly different, and the second one quantifies the increment, or decrement, of the difference D (if MK’s p-value <0.05).

All analyses were performed using the R software version 4.4.2, using the base packages as well as terra for raster data manipulation, sf (Pebesma, 2024a) and lwgeom (Pebesma, 2024b) for vector data manipulation and geometries correction, rmapshaper (Teucher and Russell, 2023) for polygons simplification, doParallel (Corporation and Weston, 2022) and foreach (Revolution Analytics and Weston, n.d.) for parallel computing, and trend (Pohlert, 2023) package for Mann-Kendall test and Sen’s slope calculations.

## Results and Discussion

By the end of 2024, PAs covered 1.7 billion hectares of semi-natural vegetation but during the 21^st^ century fire has so far burned 2 billion hectares of PAs (Fig. 1 a-d). That is, wildfires during 2001-2024 burned the equivalent of 117% of the area occupied by PAs by 2024. This does not mean that wildfires have affected the entirety of PAs, as some areas burned more than once (wildfire impacts have concentrated over 23.7% of global PA coverage, while 76.3% remains unburned). In absolute terms, the PAs most affected by fire are those in tropical and subtropical grasslands (primarily from Africa), where 1.6 billion hectares of PAs have burned, although PAs only occupy 324 million hectares (Mha) in that biome; followed by flooded grasslands, where 134.3 Mha of PAs have burned (although PAs only occupy 28.4 Mha) (Fig. 1c, d). Within forest biomes, most of protected area fire concentrates in tropical moist broadleaf forests (88.4 Mha, equivalent to 19.3% of the land covered by PAs) and in tropical dry broadleaf forests (34.2 Mha, equivalent to 82% of the PAs). The PAs least affected by fire in absolute terms were those occurring in mangroves (1.5 Mha, or 28.9% of the total area covered by PAs) and tundra (1.9 Mha, or 1.4% of the total area covered by PAs).

**Figure 1.**
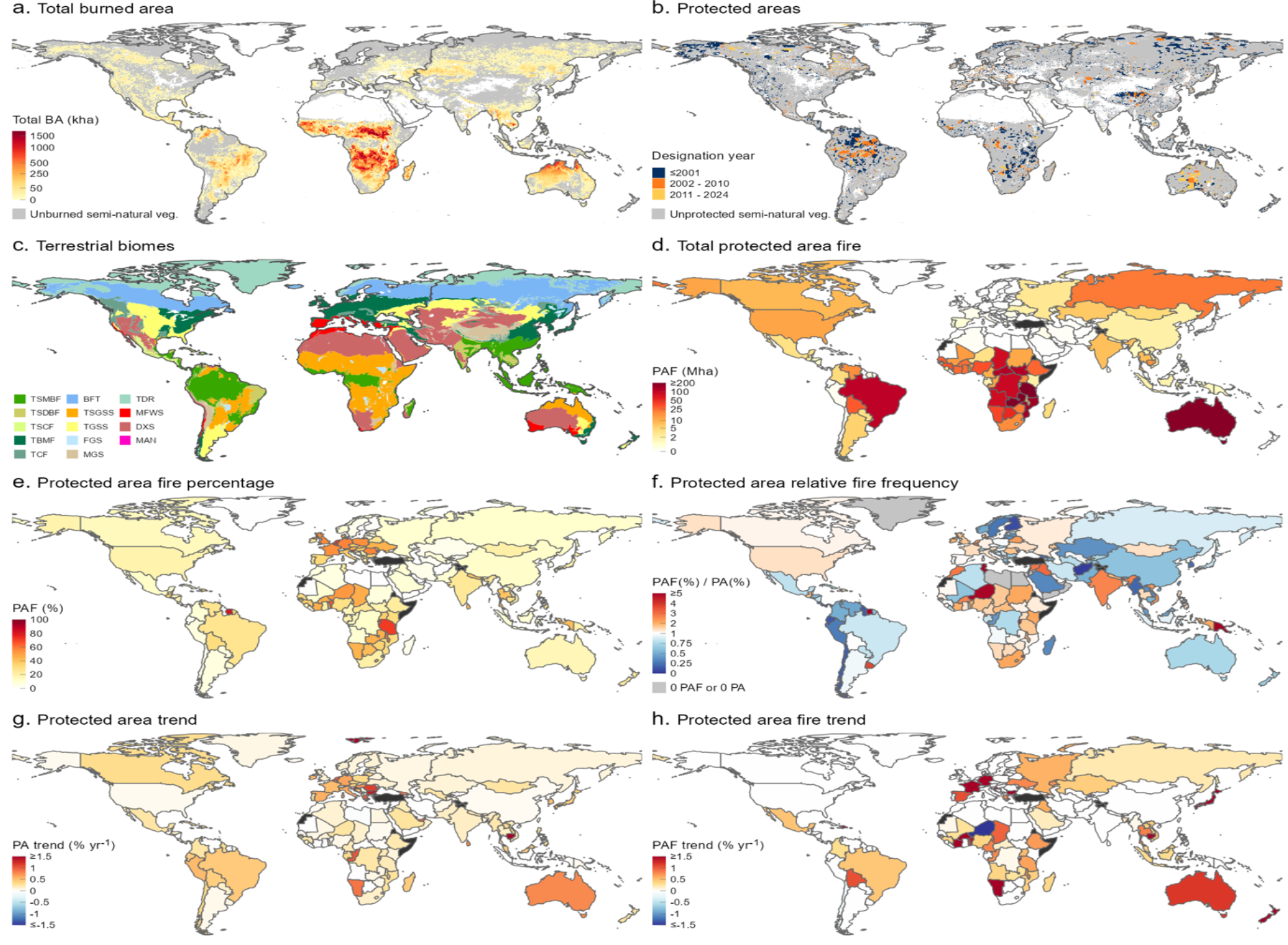
Global distribution of wildfire across protected areas during the 21^st^ century. **a-d**, Global patterns in **(a)** total area burned during 2001-2024, **(b)** protected area (PA) coverage by the end of 2024, **(c)** the distribution of biomes, **(d)** total burned area in PAs during 2001-2024, **(e)** % protected area fire (the proportion of the total burned area occurring in protected areas annually), **(f)** % protected area fire relative to the % cover of PAs. **g-h** show temporal trends in **(g)** the cover of PAs and **(h)** the % protected area fire during the 21^st^ century. Temporal trends were assessed by Sen’s slope and non-white regions indicate that the trend is statistically significant according to Mann-Kendall test (*P* < 0.05). Biome acronyms in **(c)** are: TSMBF, tropical and subtropical moist broadleaf forests; TSDBF, tropical and subtropical dry broadleaf forests; TSCF, tropical and subtropical conifer forests; TBMF, temperate broadleaf and mixed forests; TCF, temperate conifer forests; BFT, boreal forests and taiga; TSGSS, tropical and subtropical grasslands, savannas and shrublands; FSS, flooded grasslands and savannas; MGS, montane grasslands and savannas; TDR, tundra; MFWS, Mediterranean forests, woodlands and scrub; DXS, desert and xeric scrublands; MAN, mangrove. Data on protected area coverage is not available for Turkey, Eritrea, Western Sahara and the regions of Jammu and Kashmir and Azad Kashmir, which are shown as black in panels d-h.

The absolute magnitude of burned area across the different biomes, as reported above, largely reflects differences in the size and fire regimes of the different biomes. The vast majority of burned area globally occurs in tropical grasslands and savannas (specially in Africa), where fires naturally show very short rotation periods (1-5 years) (Zubkova et al., 2023). It is thus to be expected that an overwhelming proportion of the global burned area within PAs will also be concentrated in that biome. The short rotation period in tropical savanna fires (Boer et al., 2021) additionally explains why the area burned by fire is 5-fold larger than that occupied by PAs in that biome (these naturally fire-prone ecosystems were expected to burn several times over our 24 years study period).

In order to better quantify how are wildfires differentially impacting PAs across biomes, we examined the proportion of burned area occurring within protected areas (% protected area fire, or %PAF hereafter) (Fig. 1e, f) and compared it against the percent of semi-natural vegetation that is protected (Fig. 1g, h). We observed that %PAF globally increased from 18.0% in 2001 to 28.5% in 2024 (Sen’s slope annual trend size = +0.46 % yr^−1^, Mann-Kendall test, *P* <0.0001) (Fig. 2a). The increase was due partly, but not completely, to the increase in PA coverage over the same time period (from 12.6% in 2001 to 19.2% in 2024; Sen’s slope annual trend size = +0.27 % yr^−1^, Mann-Kendall test, *P* <0.0001). This result is pointing towards an overproportion in the incidence of wildfires within PAs globally, which is substantiated by a significantly larger rate of increase in %PAF, relative to the cover of PAs (Sen’s D (difference across slopes) = +0.19 % yr^−1^, Mann-Kendall test, *P*<0.0001).

**Figure 2.**
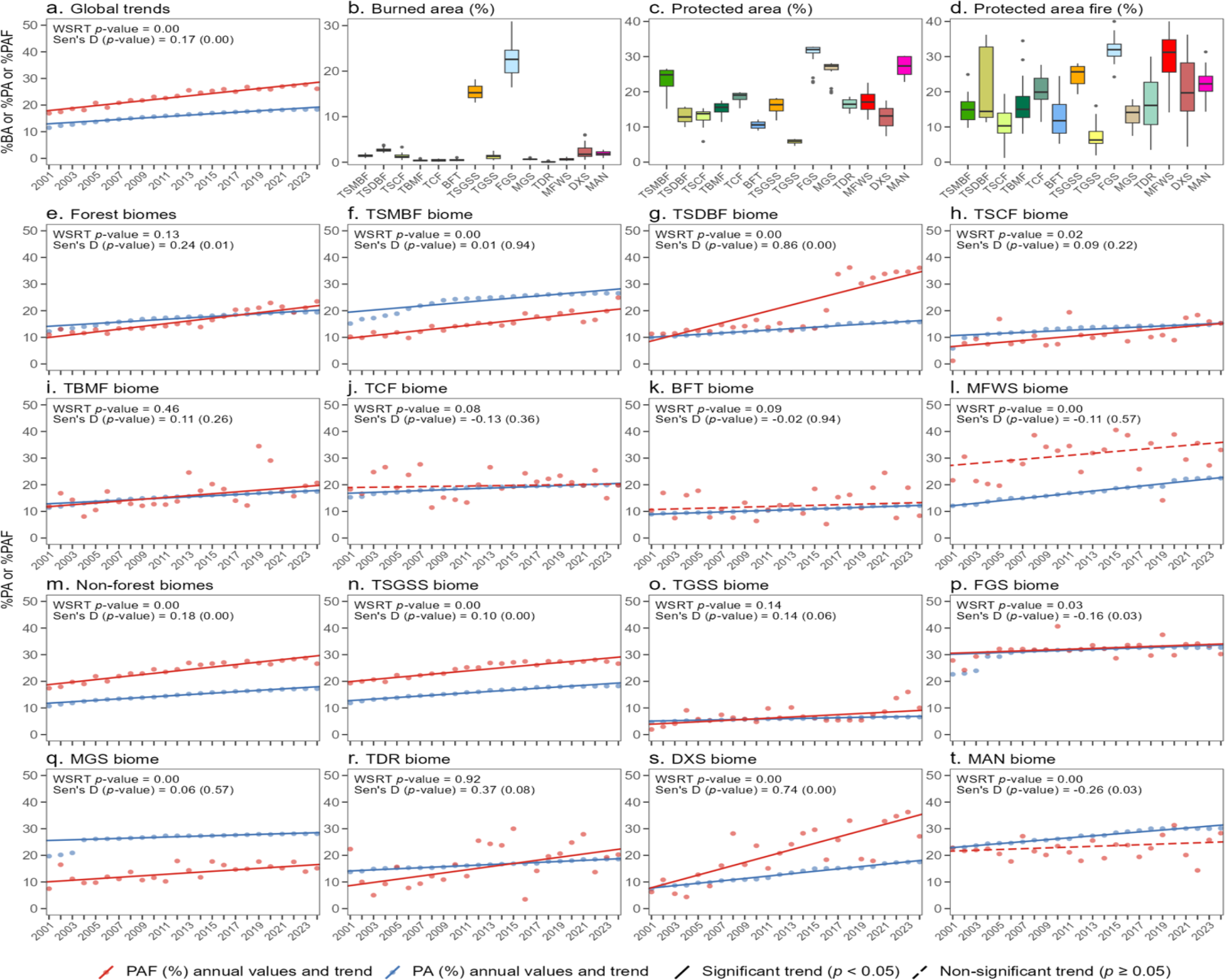
Disproportional increases in burned area have occurred globally across Protected Areas during the 21^st^ century. **(a)** Temporal trend globally in the cover of PAs (in blue) and of protected area fire (PAF, in red, the proportion of the total burned area occurring in protected areas annually). Average annual variation across biomes in **(b)** burned area, **(c)** PA and **(d)** PAF. Temporal trends in the cover of PA and PAF across forest **(e-l)** and non-forest **(m-t)** biomes. Biome acronyms are: TSMBF, tropical and subtropical moist broadleaf forests; TSDBF, tropical and subtropical dry broadleaf forests; TSCF, tropical and subtropical conifer forests; TBMF, temperate broadleaf and mixed forests; TCF, temperate conifer forests; BFT, boreal forests and taiga; TSGSS, tropical and subtropical grasslands, savannas and shrublands; FSS, flooded grasslands and savannas; MGS, montane grasslands and savannas; TDR, tundra; MFWS, Mediterranean forests, woodlands and scrub; DXS, desert and xeric scrublands; MAN, mangrove. Panels e and m refer to the overall trends for all forest and all non-forest biomes, respectively. The lines indicate the results of Sen’s slope analyses. WSRT indicates Wilcoxon Signed Rank Test, with significant differences indicating differences in mean values across the time-series. Sen’s D refers to the difference in % across both slopes (PAF – PA) and the p-value indicates whether slopes are significantly different according to a Mann-Kendall test.

This overproportion was consistent across PAs in forest and non-forest biomes overall (Figs. 2 a,b), but it was not universal and major differences arose across biomes (Figs. 2c-p). Among forest biomes, %PAF averaged over the study period was significantly higher than the cover of PAs in Mediterranean (+12.5%, Wilcoxon test *P* > 0.0001) and tropical dry (+7.4%, Wilcoxon test *P* > 0.0001) forests, and it was marginally higher in boreal (+2.0%, Wilcoxon test *P* = 0.09) and temperate (+1.8%, Wilcoxon test *P* = 0.08) conifer forests. In contrast, in tropical moist broadleaf and conifer forests, %PAF was significantly lower (−8%, Wilcoxon test *P* < 0.0001; and −1.8%, Wilcoxon test *P* = 0.02, respectively) than the cover of PAs, indicating that PAs in these tropical biomes suffer a disproportionately lower impact of wildfires than that expected from their coverage. Finally, there were no differences between %PA cover and %PAF in temperate broadleaf forests. The disproportionate impact of wildfire over protected areas is thus mostly limited to drier and hotter forests, and only marginally apparent in non-tropical conifer forests.

In non-forest biomes, we similarly observed diverging impacts of wildfire in PAs. An overproportion in protected area fire has occurred in tropical (+8.7%, Wilcoxon test *P* < 0.0001) and flooded grasslands (+ 1.2% Wilcoxon test *P* = 0.031), and also in desert and xeric shrublands (+7.8%, Wilcoxon test *P* < 0.0001). Conversely, PAs burned at significantly lower proportions than expected in montane grasslands (−12.7%; Wilcoxon test, *P* < 0.0001) and mangroves (−4.7%, Wilcoxon test *P* < 0.0001). In the remaining biomes (temperate grasslands and tundra), %PAF was consistent with the cover of PAs. The disproportionate effect of %PAF within non-forest ecosystems seems to be limited to those biomes that experience seasonal droughts.

The observation that an overproportion of protected area fire tends to occur in seasonally dry, fire-prone biomes (e.g. tropical grasslands or Mediterranean forests), while the opposite pattern (underproportion of protected area fire) is more prevalent where fires are putatively rare (e.g. tropical moist broadleaf forests (Boer et al., 2021)), deserves further attention. This pattern could arise from two different processes: the first is that the establishment of protected areas impacts on the fire regime. Alternatively, it is also possible that areas that are now protected already had higher/lower fire activity before protection was established. Our results could additionally reflect the impacts of the exclusion of Indigenous and cultural burns from “wilderness” areas (Mariani et al., 2024; Minnich, 1983). The goal of this study was to assess and quantify the biogeographical pattern of fire activity on PAs, as no study had yet attempted this globally and previous regional studies revealed differing results. Future research efforts should thus address these hypotheses, to better understand the processes underlying the patterns we document here, given their importance for a successful GBF implementation. Regardless of the mechanism, our results do highlight the need to consider fire management in PAs.

The overproportion of fire in PAs from fire-prone biomes should not be necessarily considered a negative outcome, as wildfires play a key role in the health of those biomes (Moritz et al., 2014). However, extreme wildfires have become more common over the last few years (Cunningham et al., 2024). These extreme events may hinder recovery if increases in fire intensity, frequency, severity or area burned push the fire regime beyond the species realized niches (Doherty et al., 2024; Nolan et al., 2021).

Beyond ecological impacts, wildfires in PAs can also affect infrastructure and people and, in those cases, land management for human safety (including ignitions control and fuel management) should take priority. If the current disproportion between %PAF and PA cover continues (e.g. 28.5% PAF vs 19.2% PAs cover in 2024), we can expect that 44.5% of the area burned globally will be concentrated within PAs, if PA coverage reaches GBF’s 30% target in semi-natural ecosystems. Under an increasing climate emergency, we can expect further increases in the disproportionate incidence of wildfires in PAs, potentially compromising the goals of the Global Biodiversity Framework, and other conservation efforts in some of the world’s most ecologically valuable ecosystems.

## Acknowledgements

We acknowledge funding from the Spanish MICINN (PID2022-138158OB-I00) the European Union’s Horizon 2020 research and innovation programme under grant agreement no. 101003890 project FirEUrisk, the National Science Foundation of China project U20A2079, the Sichuan Government 2024YFFK0405 and the Westpac Fellowship Program.

## Author contributions

VRD designed the study, obtained funding and drafted the manuscript. ACC conducted the research. ACC, HC, YH, OZ, HY, RD and YY assisted with data analyses, interpretation and contributed to revisions.

